# Validation of an MPS based intestinal cell culture model for the evaluation of drug-induced toxicity

**DOI:** 10.1101/2024.07.18.604106

**Authors:** Stefanie Hoffmann, Philip Hewitt, Isabel Koscielski, Dorota Kurek, Wouter Strijker, Kinga Kosim

**Affiliations:** Early Investigative Toxicology, Chemical and Preclinical Safety, Merck Healthcare KGaA, Darmstadt, Germany; MIMETAS B.V., Oegstgeest, The Netherlands

**Keywords:** OrganoTEER, Organ-on-chips, in vitro barrier model, drug toxicity, Caco-2, pre-clinical safety, gastrointestinal-toxicity

## Abstract

The potential for drug-induced gastrointestinal (GI) toxicity is significant, since the GI tract is one of the first barriers which come in to contact with oral drugs. In pharmaceutical research, the complex behavior of human intestinal cells is traditionally investigated using 2D cultures, in which one cell type grows under static conditions. With the development of advanced microphysiological systems (MPS) more in vivo like conditions can be generated which increase the physiological nature and also the predictive validity of these models.

Caco-2 cells are known for their capability to build tight junctions. These connections are responsible for the maintenance of intestinal homeostasis and can be used as a specific safety endpoint, by measuring the Trans Epithelial Electrical Resistance (TEER), for the investigation of drug-induced GI toxicity.

Compared to a widely used Caco-2 cell 2D Transwell model, the advanced MPS model (Mimetas OrganoPlate®) allows for the recapitulation of the enterocyte cell layer of the intestinal barrier as the Caco-2 cells grow in a tubular structure through which the medium continuously flows.

The OrganoPlate® intestinal model was qualified to be implemented as a routine test system for the early prediction of drug-induced GI toxicity based on the measurement of the tightness of the cell layer by measuring changes in the TEER. For this qualification 23 well known compounds as well as a positive, negative and solvent control were selected. The compounds were selected based on their known effect on the GI system (chemotherapeutics, tight junction disruptor, liver toxins, controls, NSAIDs and a mixed group of drugs).

The TEER values were measured 4h and 24h after treatment. In parallel the cell viability was determined after 24h to be able to distinguish between an unspecific cytotoxic effect or direct tight junction damage. Overall, from the 23 tested compounds, 15 showed the expected outcome, i.e.,the compound led to a decrease of the TEER for the positive control compounds, or the TEER value remained stable after treatment with non-GI-toxic compounds. In summary, this MPS model allowed the recapitulation of the human intestinal GI barrier and will enable a faster and more robust assessment of drug-induced damage in the GI tract.

## Introduction

Since the gastrointestinal (GI) tract with its mucus layer is one of the first barrier that comes into contact with external contaminants, toxins, foreign substances and food, it has an important protective function (Paone and Cani (2020). A single layer of intestinal epithelial cells (IECs) organized into crypts and villi builds the mucosal surface of the gastrointestinal tract. This monolayer consists of several cell types which differentiate from epithelial stem cells (Soderholm and Pedicord 2019). These cells separate the gut lumen from the blood and act as barrier against xenobiotics, while allowing the absorption of nutrients (Vancamelbeke and Vermeire 2017). The tight barrier is mainly generated by the presence of polarized absorptive enterocytes connected via different junctional complexes. These connections are divided in tight junctions, adherens junctions, desmosomes, and gap junctions (Turner 2009).

Damage to the tight junctional complexes increases the paracellular permeability and can lead to tissue damage, and eventually inflammation (Lee 2015). This increased permeability can be detected by using a quantitative non-invasive, label free, real-time measurement with an impedance-based instrument (Benson, Cramer et al. 2013). This electrical measurement (Trans Epithelial Electrical Resistance (TEER)) is the gold standard for monitoring the barrier function of epithelial cells in vitro. To determine the efficacy and toxicity of new drug candidates, as early as possible typically in vitro models are used.

The cultivation of cells on semipermeable membranes enables the generation of two compartments. An electrode is placed in each of the compartments to allow ohmic resistance measurement (Srinivasan, Kolli et al. 2015, Nicolas 2021). The resulting TEER values are reliable indicators of the barrier integrity.

A classical and widely used method is the measurement of the TEER of a 2D cell monolayer of Caco-2 cells cultured on semipermeable membranes. Caco-2 cells are originally from a human colon adenocarcinoma but express functional and morphological characteristics of differentiated small intestinal enterocytes. They form confluent layers with highly polarized cells, joined via tight junctions, including apical and basolateral sides with microvilli on the apical membrane (Sambuy, De Angelis et al. 2005). The full differentiation of Caco-2 cells takes around 14-21 days on Transwell inserts and reaches TEER values of 150-400 Ω (Srinivasan, Kolli et al. 2015).

Overall, there are multiple commercial cell culture models available which mimic the intestinal barrier (Table 1).

**Table 1:** Overview on commercially available cell culture models for the intestinal tract (non-exhausive)

|  | Model |  |  |  |  |  |  |  |
| --- | --- | --- | --- | --- | --- | --- | --- | --- |
|  | Caco-2 | Caco-2 & HT29 | Organoids | RepliGut® | Mimetas OrganoPlate® | EpiIntestinal™ 3D | Ussing chamber | Animals |
|  | In vitro | In vitro | In vitro | In vitro | In vitro | In vitro | Ex vivo | In vivo |
| Source | Human, cancer cell line | Human, cancer cell lines | Human, iPSC or adult stem cells | Human, primary | Human, cancer cell line | Human, small intestine epithelial and endothelial cells | Animal tissue explant | Complete animal |
| Cell types | Monoculture of Caco-2 | Co-culture of Caco-2 & HT29-MTX | Multiple cells (primary or adults/iPS cells) | Multiple cells (colon or small intestine) differentiated from stem cells | Monoculture of Caco-2 | Multiple (Enterocytes, paneth cells, tuft cells, M cells, stem cells) | Multiple species | All cells from an animal |
| Platform | Transwell | Transwell | Flat plate | Transwell | 3lane OrganoPlate | Transwell | - | - |
| Flow | no | no | no | no | yes | no | no | yes |
| ECM | no | no | yes (Matrigel) | no | yes (Collagen) | yes | no | yes |
| Pre-preparation time* | 21 days | 21 days | ~6-10 days | 14-17 | 4-6 days | Up to 14d (MatTek Corp.) | Animal growth | Animal growth |
| Pro's | - cheap<br>- easy handling | -cheap<br>- easy handling | - 3D<br>- Presence of important cell types from the tissue | Proliferative stem cells | - flow introduce sheer stress and nutrient transport<br>-compatible with commercial plate reader | -forms villi, microvilli, brush border and tight junctions<br>-cultivation up to 42d<br>-organotypic small intestinal tissue reconstruct | - barrier of animal tissue | - Complex structure<br>- cell- cell contact and organ interaction<br>- complete organism (DNA) |
| Con's | - cancer cell line<br>- 2D<br>- | - cancer cell line<br>- 2D | - high deviations | - needs training | - needs training | - needs training | - needs animals<br>- analysis of a small set of segments of epithelial tissue<br>- low throughput | - Animal experiment<br>- Expensive<br>- Time intensive<br>- ethical reasons<br>-Lack of applicability |
| Supplier | ATCC & others | ATCC & others | several | Altis biosystem | Mimetas | Mattek | Reprocell & others | several |
| Costs | \$ | \$ | \$\$ | \$\$\$\$ | \$\$ | \$\$\$ | \$\$- \$\$\$ | \$\$\$\$\$ |
| Reference | (Sambuy, De Angelis et al. 2005) | (Walter, Janich et al. 1996) | (Zhang, Huang et al. 2021) | (Lemmens, Van Camp et al. 2021) | (Trietsch, Naumovska et al. 2017, Nicolas 2021) | (Ayehunie 2013) | (Thomson, Smart et al. 2019) | (Crystal 2018) |

\*Pre-preparation time = time for pre-cultivation of animals/cells for experiments

None of them can completely cover the complex structure of the human intestine but each has its benefits and limitations. Micro physiological systems (MPS) and Organ-on-a-chip (OoC) systems allow the combination of several parameters, such as extracellular matrices (ECM), three dimensional (3D) growing of cells, nutrient flow including shear stress, and co-culture options. By an improved mapping of the in vivo environment, 3D models can help to better select drug candidates and can reduce costs by improving the prediction of drug efficacy and toxicity (Peng, Datta et al. 2017).

A good combination of complexity and throughput can be created with the Mimetas OrganoPlate® 3-lane, and therefore chosen for this study. This system includes cells which grow against an ECM in a 3D tubular structure. The nutrient supply is generated by a bidirectional flow of media.

## Methods

### Cell culture and seeding of cells

The human colon carcinoma cell line Caco-2 (Sigma-Aldrich, 86010202) was cultured in T175 flasks in DMEM 4.5g/L glucose (Lonza, 12-614F), 10% FBS (Gibco, 102070-106), 1% Penicillin /Streptomycin (Sigma, P4333) and 1% L-Glutamine (Gibco, 25030-081). Caco-2 cells were harvested between passage 3 and 20 for experiments. The cells were cultured at 37°C and 5% CO2 and every 2-3 days medium were refreshed. At 70-80% confluency the cells were either sub-cultured or used for experiments.

Cells were trypsinized using Trypsin-EDTA solution (Sigma-Aldrich, T3924) and resuspended at 10000 cells per µl prior to seeding. The cells were applied to 3-lane OrganoPlates (Mimetas, 4004-400-B). Prior cell seeding 1.7µl of gel composed of 1M Hepes (Gibco, 15630-080)), 37g/L NaHCO3 (Sigma, S5761) and 4mg/ml Collagen I (R&D System, 3447-020-01) in a ratio 1:1:8 was added in the gel inlet and incubated for 15min at 37°C. After polymerization, 50µl HBSS (Cytiva, SH30268.01) was dispensed in the gel inlet channels to avoid dehydration of the gel. The cells were applied by seeding 2µl of 1x 107 cells/ml cell suspension in the inlet of the top channel. Subsequently, 50µl of medium was added to the same well. Afterwards the OrganoPlate was placed on its side on a specific plate stand for 4h at 37°C to allow the cells to settle against the ECM. This was followed by adding additional 50µl medium to the outlets of the top channel and to the inlets and outlets of the bottom channel. Next, the OrganoPlate was placed horizontally in an incubator (37°C and 5% CO2) on an interval rocker (OrganoFlow, Mimetas), which switches the inclination between +7° and −7° every 8min which allows a bidirectional flow of the medium. Every 2-3 days the medium was refreshed.

The 3-lane OrganoPlate® contains 40 chips of which each has three microfluidic channels, and all is based on a 384well plate format. Each chip contains of two perfusion channels and one channel for an extracellular matrix (Figure 1). Compared to commonly used 2D models it has a shorter pre-cultivation time (2D Transwell: 14-21 days until polarized Caco-2 cells and OrganoPlate: 4-6 days), more complexity (3D tube of cells, ECM, media flow) and it allows the measurement of the TEER of 40 chips in parallel.

**Figure 1:**
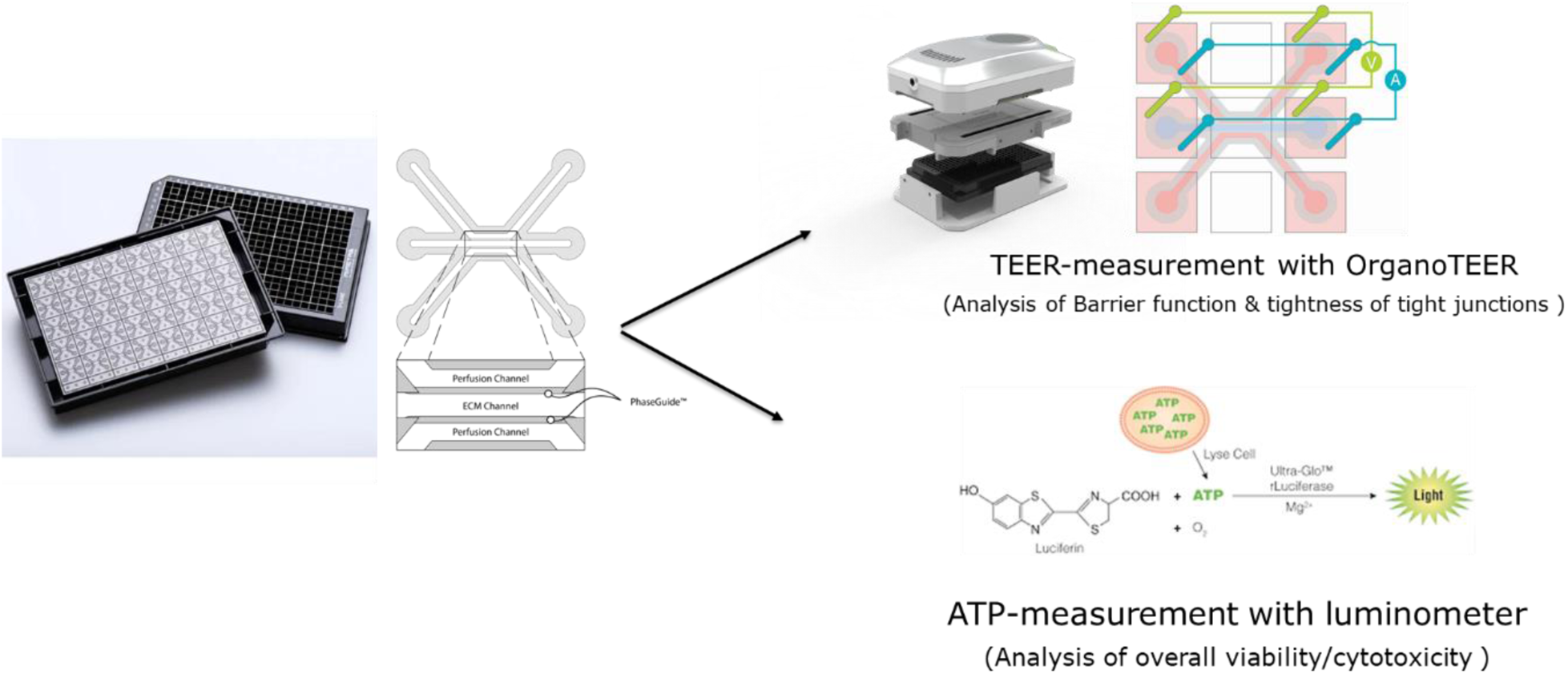
Schematic overview of the OrganoPlate 3-lane and the performed assays in this study

### Treatment of cells

Caco-2 cells in OrganoPlates were exposed to test compounds (Table 2) for 24h. For each compound three different concentrations were tested. For the treatment the medium was completely removed and 50µl of compound-medium solution was added in each inlet and outlet of the top channel. Compounds were selected based on the occurrence of side effects in the GI tract or based on the occurrence of tight junction damage specifically. Compound concentrations were chosen based on literature values and already published data with Caco-2 cells in 2D. Each compound was tested three times.

**Table 2:** Reference compounds which were used for testing the usability of the OrganoPlate to predict well-known GI toxicity.

| Compound | Manufacturer | Catalogue number |
| --- | --- | --- |
| Ibuprofen | Cayman Chemicals | 70280 |
| Indomethacin | Sigma Aldrich | I7378 |
| Diclofenac | Sigma Aldrich | SML3086 |
| 5-Fluorouacil | Sigma Aldrich | F6627 |
| Alosetron | Cayman Chemicals | 22434 |
| Irinotecan hydrochloride | abcr | AB252173 |
| Sunitinib | Sigma Aldrich | PZ0012 |
| Sorafenib | Sigma Aldrich | SML2653 |
| Flavopiridol | Cayman Chemicals | 10009197 |
| Bortezomib | VWR | 5.04314.0001 |
| Bosutinib | Sigma Aldrich | PZ019 |
| Carboplatin | Cayman Chemicals | Cay13112 |
| Docetaxel | Cayman Chemicals | Cay11367 |
| Afatinib | LKT Laboratories | LKT-A2077.5 |
| Troglitazone | Cayman Chemicals | 71750 |
| Trovaflaxacin | Sigma Aldrich | PZ0015 |
| Homoharringtonin | Sigma Aldrich | SML1091 |
| Patulin | VWR | 5.04314.0001 |
| Phenylarsine oxide | VWR | A14674.03 |
| Sodium orthovanadate | Sigma Aldrich | 567540 |
| Duloxetine | Sigma Aldrich | SML0474 |
| Terfenadine | Sigma Aldrich | T9652 |
| Loperamide hydrochloride | LKT Laboratories | L5660 |
| Staurosporine | abcr | AB353975 |
| Metformin hydrochloride | Sigma Aldrich | PHR1084 |

### TEER measurement

The transepithelial electrical resistance (TEER) of the Caco-2 cultures in the OrganoPlate was measured by using an automated multichannel impedance spectrometer (OrganoTEER, Mimetas). TEER was measured before treatment (0h), 4h after treatment (4h) and 24h after treatment (24h). Before the measurement, a medium exchange of 50µl medium was performed in the inlets and outlets of the middle channel and then the OrganoPlate was equilibrated for 30min at RT. The electrode board of the OrganoTEER perfectly fits in the OrganoPlate and introduce the electrodes in the medium of all inlet and outlet wells connecting to the apical and basal side of the Caco-2 tubes.

Before each measurement the electrode board was cleaned by spraying 70% ethanol onto the board. Under a laminar flow the electrode was left for drying for at least 30min. Between each experimental run the electrode board was immersed in a single well plate filled with 50ml of a 1:20 solution of RBS T342 (Chemical Products R. Borghgraef N.V, BE) in Milli-Q-Water. The electrode board was left in the solution for 15-20min and afterwards the electrode board was rinsed with Milli-Q-Water and left to dry ambiently.

### ATP measurement

To quantify the number of viable cells the CellTiter-Glo® 3D (Promega, G9681) assay was used, which determines the amount of intracellular ATP. And is indicative of the number of metabolically active cells. 25µl CellTiter-Glo® 3D reagent was added in each inlet and outlet of the top channel. The OrganoPlates were placed on a shaker for 5min, 300rpm in the dark and then left for 25min. Immediately the luminescence signal was read in a luminometer.

### Data analysis and statistics

Data from the OrganoTEER was analyzed using the OrganoTEER software and excel for the calculation of the remaining TEER (%) values. The remaining TEER (%) values are indicating the TEER values after the measurement in % in relation to the start measurement (TEER measurement before the treatment).

## Results

The selected compounds were divided into six subclasses (controls, NSAIDs, chemotherapeutics, liver toxins, tight junction disruptor and a mixed group of compounds) according to their reported effect (Table 3). Compounds were selected based on their effect in vivo or on Caco-2 cells. Compound concentrations were selected based on given concentrations, which showed toxic effects, from the literature.

**Table 3:** Overview of the selected reference compounds used for the qualification of the OrganoPlate and the use of the OrganoTEER to predict GI toxicity in vitro.

| Compound class | Compound name | MoA / toxicity | Side effects in the human GI tract | Reference |
| --- | --- | --- | --- | --- |
| Controls | Metformin | Type 2 diabetes mellitus drug; lowers glucose production in the liver and increase glucose utilization in the gut | In adequate amounts: Diarrhea, nausea, vomiting, abdominal discomfort | (Bonnet and Scheen 2017, Rena, Hardie et al. 2017, Peters, Landry et al. 2019) |
|  | Staurosporine | Strong protein kinase and cyclin dependent kinase inhibitor, induces cell cycle arrest | No side effects known for the GI tract | (Oikawa, Shimamura et al. 1992, ANTONSSON and PERSSON 2009, Trietsch, Naumovska et al. 2017) |
| NSAID | Ibuprofen | Inhibition of prostaglandin synthesis, antibody synthesis, cytokine production, transcription factors and inhibition of leucocyte functions; apoptosis, | Abdominal pain, nausea, diarrhea | (Rampal, Moore et al. 2002, Rainsford 2009) |
|  | Diclofenac | Inhibition of prostaglandin synthesis COX-inhibitor | Bleeding, inflammation, and ulceration in SI; nausea, vomiting, diarrhea, dyspepsia, abdominal pain; rare: hemorrhage, gastrointestinal ulcer, colitis, constipation | (Bhatt, Gunasekara et al. 2018) |
|  | Indomethacin | Inhibit prostaglandin synthesis, COX-inhibitor | Bleeding, ulceration, perforation of stomach or intestine, lesions in the small intestine | (Maseda and Ricciotti 2020) |
| Chemotherapeutics | 5-Fluorouracil | toxic effect on specific proteins of the tight junctions in immunodeficient mice, specifically on occludin and claudin-1 Both proteins are also found in the human intestine and have also been detected in Caco-2 cells; cell death | Intestinal injury, epithelial ulceration in the mucosa (mucositis) which manifests mainly in pain and dyspeptic syndromes or inflammation; damage of intestinal barrier function, reduced enterocyte proliferation and crypt cell apoptosis | (Soares, Mota et al. 2013, Song, Park et al. 2013, Garcia-Hernandez, Quiros et al. 2017) |
|  | Alosetron | highly potent 5-HT<sub>3</sub> receptor antagonist and improves abdominal pain and can slow down colonic transit | Constipation, ischaemic colitis, and death | (Peters, Landry et al. 2019) |
|  | Irinotecan | Accumulation of SN-38 metabolite in the enterocytes; SN-38 inhibits the DNA topoisomerase I by inducing DNA damage in tumor cells and accumulation in intestinal mucosa | Nausea, vomiting, abdominal pain, constipation | (Lee, Ryan et al. 2014, Wardill, Bowen et al. 2014, EMC 2018) |
|  | Sunitinib | Tyrosine kinase inhibitor; used for the treatment of renal carcinoma and gastrointestinal stromal tumors | Nausea, vomiting, indigestion | (Le Tourneau, Raymond et al. 2007, Durand 2012, Sehdev 2016) |
|  | Sorafenib | Tyrosine kinase inhibitor; | Diarrhea; decrease in TEER and plasma citrulline (Caco-2) | (Durand 2012) |
|  | Flavopiridol | Inhibits cyclin dependent kinases and blocks cell cycle progression | Diarrhea, nausea, vomiting | (Senderowicz 1999, Byrd, Lin et al. 2007, William, Mitch et al. 2010) |
|  | Bortezomib | Proteasome inhibitor | Nausea, Vomiting, Constipation, Diarrhea | (Sun, Wilkins et al. 2005, Peters, Landry et al. 2019) |
|  | Bosutinib | Tyrosine kinase inhibitor, Decrease TEER | Nausea, Vomiting, bleeding | (Khouri, Cortes et al. 2012, French 2019, Peters, Landry et al. 2019) |
|  | Carboplatin | Apoptosis inducer, DNA crosslinker | Nausea, Vomiting | (Peters, Landry et al. 2019) |
|  | Docetaxel | Taxane, disrupts the normal function of microtubules | Nausea, Vomiting, enterocolitis, perforation, colitis, diarrhea | (Ho and Mackey 2014, Peters, Landry et al. 2019) |
|  | Afatinib | EGFR inhibitor | Diarrhea | (Peters, Landry et al. 2019) |
| Liver toxins | Troglitazone | Liver tox | No side effects known for the GI tract | - |
|  | Trovaflaxacin | Liver tox | No side effects known for the GI tract | - |
| Tight junction damager | Phenylarsine oxide | PTP inhibitor, which is a key mediator of intestinal epithelial barrier function. Caco-2: downregulation of ZO-1, disruption of claudins leads to tight junction; decrease in TEER | In humans: decrease in surface area of the GI tract, change in TEER | (Rao, Baker et al. 1997, Mahfoud, Maresca et al. 2002) |
|  | Sodium orthovanadate | Inhibitor of protein tyrosine phosphatases, alkaline phosphatase and ATPase, disrupt the tight junctions | Increases intestina, epithelial permeability and changes homeostasis | (Rao, Baker et al. 1997, Kim, Jeong et al. 2022) |
|  | Homoharringtonine | Inhibits protein synthesis (ribosomes) of cancer cells | Increases intestinal epithelial permeability (Caco-2), reduces tight junction barrier (Caco-2) | (Zhou, Zittoun et al. 1995, Watari, Hashegawa et al. 2015) |
|  | Patulin | Mycotoxin, alters intestinal barrier function | induces intestinal injuries, including epithelial cell degeneration, inflammation, ulceration, and hemorrhages; decrease TEER (Caco-2) | (Mahfoud, Maresca et al. 2002) |
| Mixed group | Duloxetine | inhibits the reuptake of serotonin and norepinephrine (NE) in the central nervous system | Nausea, vomiting, constipation | (Peters, Landry et al. 2019) |
|  | Loperamide | Opiate agonist, inhibits action of calmodulin | Bloating, nausea, vomiting, constipation | (Mellstrand 1987, Hanauer 2008) |
|  | Terfenadine | prolongation of the QT interval | Nonspecific | (Roden 2019) |

In prior experiments (data not shown) the OrganoPlates® was shown to generate a stable TEER value after 6 days. For an internal validation of the OrganoPlate and the use of the OrganoTEER for the measurement of the tightness of the cell barriers, 23 well-known compounds were used to treat the Caco-2 cells.

### Characterization of Caco-2 cells in the OrganoPlate

After seeding of the Caco-2 cells in the top channel inlet the cells start to grow against the extracellular matrix which is made of collagen I. The cells grew in a tubular structure within 4 days (Figure 2) and formed a tight monolayer. The staining with the antibodies Claudin 7 and E-cadherin indicates an undisrupted appearance of tight junctions, and this is correlated to a tight barrier function (Figure 3).

**Figure 2:**
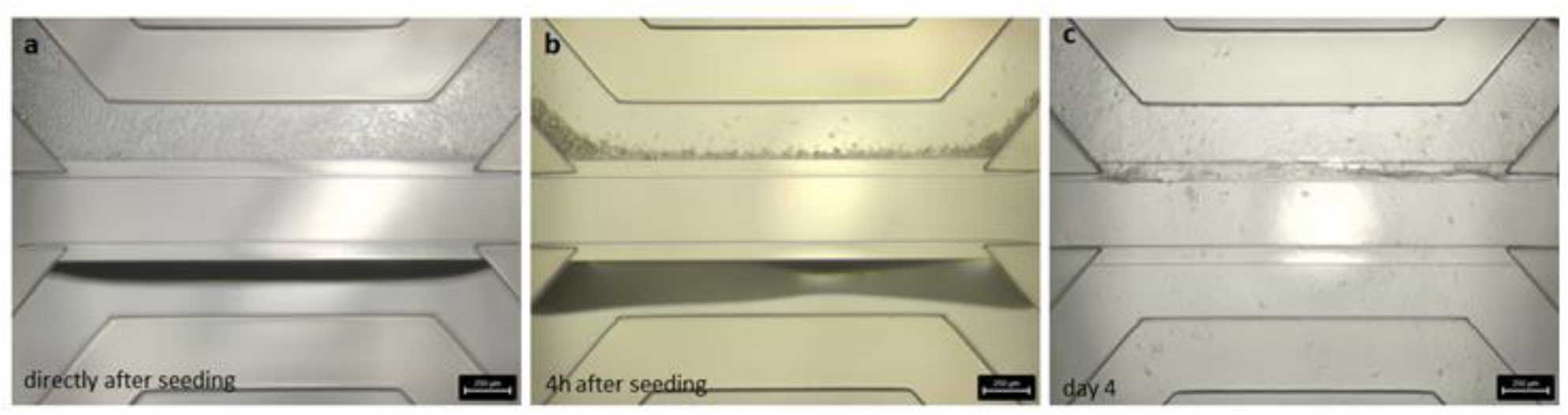
Brightfield images of the seeded Caco-2 cells in the OrganoPlate® 3-lane

**Figure 3:**
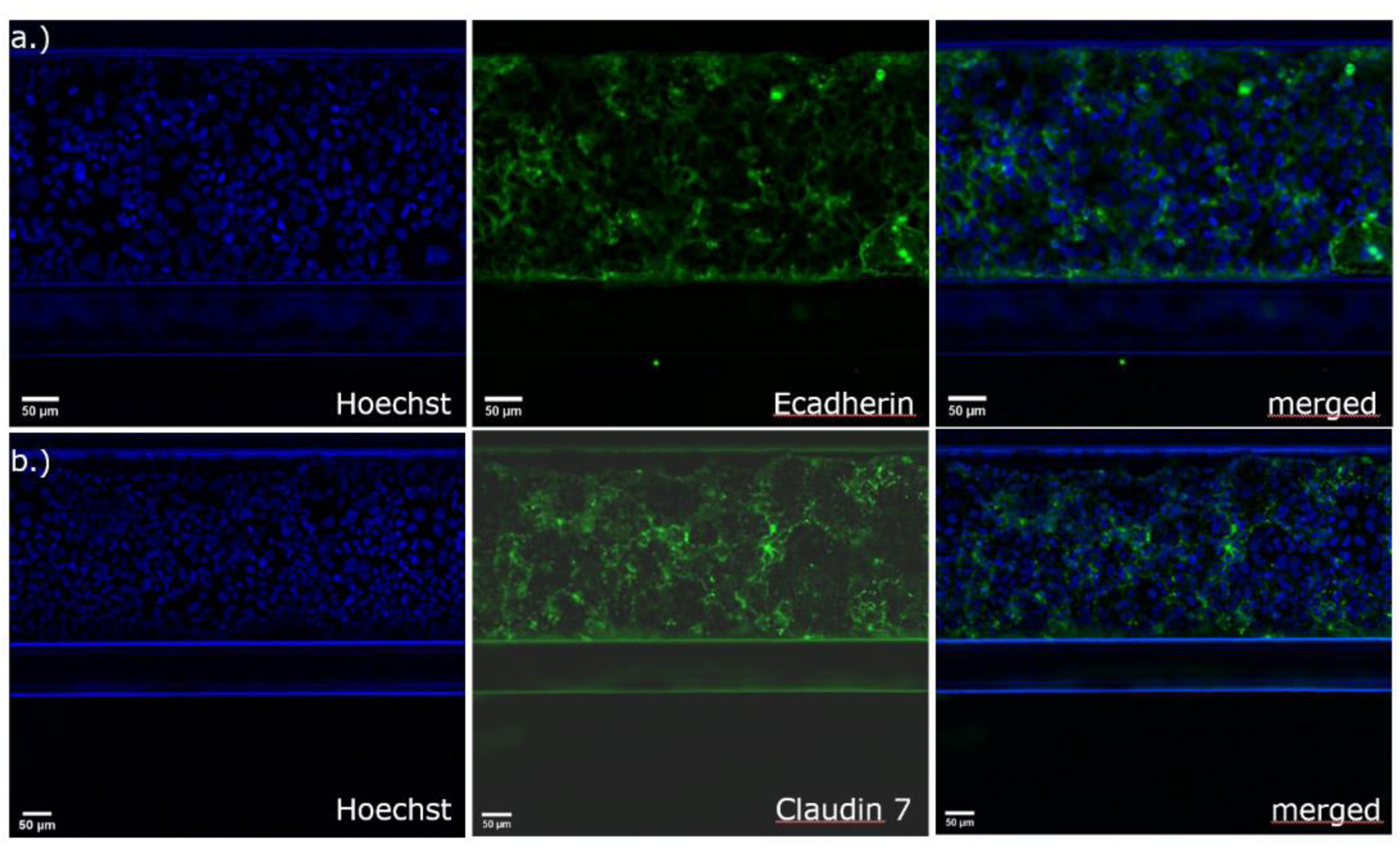
Immunofluorescence staining with a.) E-cadherin for the detection of adherens junctions and b.) claudin-7 for the detection of tight junctional complexes in Caco-2 cells in the 3-lane OrganoPlate®

### TEER measurements after test compound exposure

We assessed the barrier function of Caco-2 tubes in response to the control compounds (staurosporine and metformin) and to the test compounds which are listed above (Table 2). Staurosporine, a protein kinase C inhibitor which induces apoptosis, is known to disrupt the epithelial barrier, and was selected as a positive control and metformin, which is used for the treatment of type 2 diabetes was used as negative control.

The positive control staurosporine lead to a reduction of the TEER value already 4 h after treatment at all tested concentrations (100, 10, 1 µM) but with a stronger reduced TEER after 24 h of treatment (Figure 4a). The viability results showed a dose dependent decrease of viability after 24 h after treatment with staurosporine (Figure 4b). Compared to this, the TEER values for metformin and DMSO remained stable after 4 h and 24 h but also the viability of the cells remained at approximately 100 % after 24h (Figure 4a and 4b).

**Figure 4:**
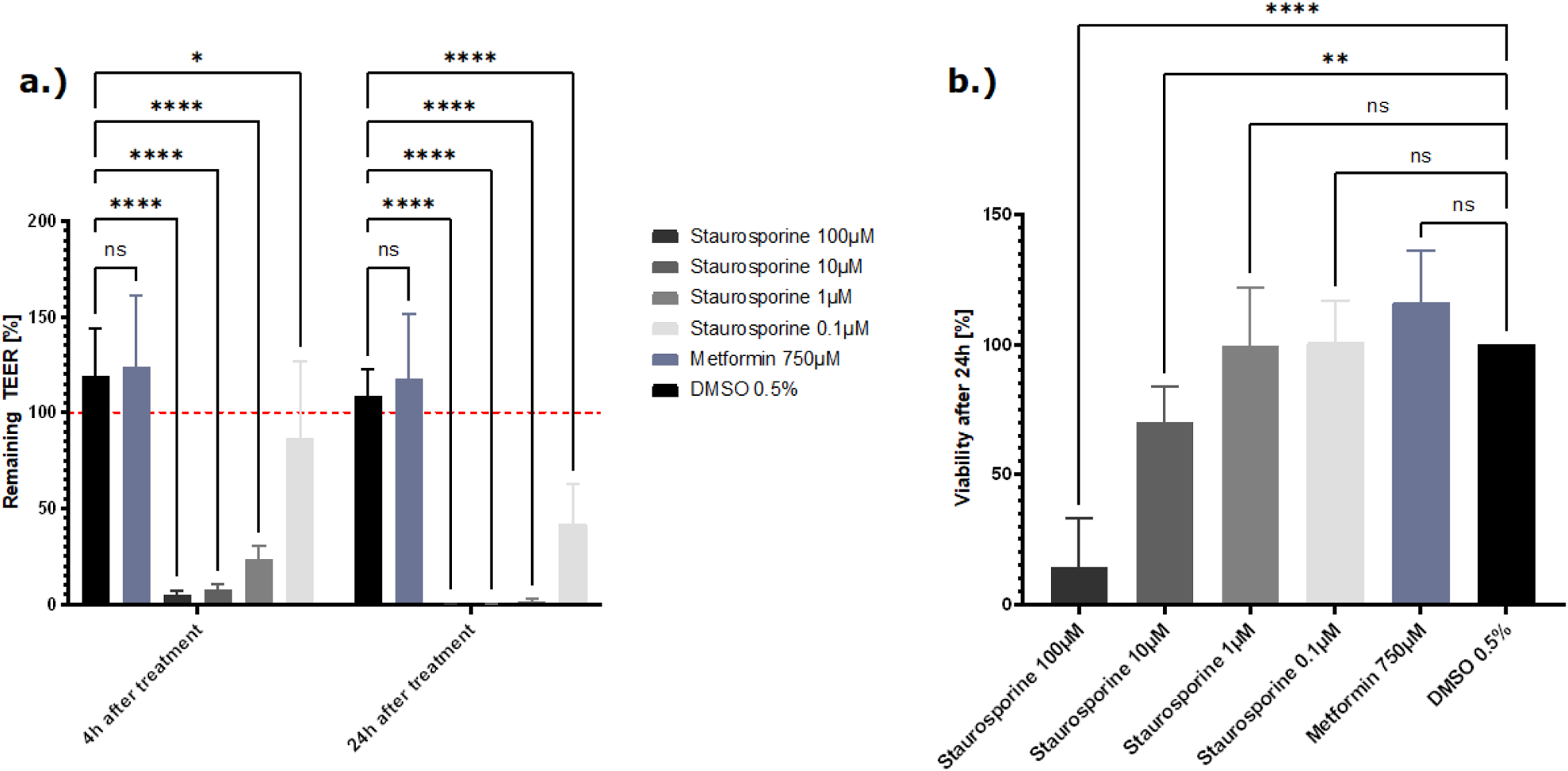
a.) Effect of the control compounds on the barrier function represented as % TEER to the t0 of the treatment. Measured TEER values after treatment with the positive control staurosporine, vehicle control DMSO and the negative control metformin. Shown are the mean remaining TEER values 4h and 24h after treatment. The red dotted line shows the normalized start TEER values of 100%. b.) Effect of the control compounds on the cell viability shown as % change to the DMSO control. Measured viability with the CellTiter Glo 3D kit, 24h after treatment with staurosporine, DMSO and metformin. The data are presented as means ± SD (n=8, statistical analysis of TEER values: **** p<0.0001 and ** p=0.0013 by two-way ANOVA with Dunnett’s test for multiple comparisons to the control. For statistical analysis of Viability values: ****p < 0.0001 and ***p=0.0037 by one way ANOVA with Dunnett’s test for multiple comparisons to the control).

The reference compounds were tested at least in 3 concentrations which were selected based on already published data or previous internal experiments. Based on their field of application the compounds were subdivided in six compound classes (NSAIDs, chemotherapeutics, tight junction disruptors, a mixed group, liver toxins and control compound).

From the NSAIDs tested, ibuprofen did not influence the tight junctions or the viability of the cells. Treatment with indomethacin leads to a decrease of the TEER at the highest concentration tested (500 µM) without a decrease of viability (Supplementary Figure 1). Diclofenac damages the tight junctions and reduces the viability after treatment with the two highest concentrations (2000 and 1000 µM) (Figure 5). The results for the TEER and viability experiments clearly show a dose dependent toxicity of diclofenac.

**Figure 5:**
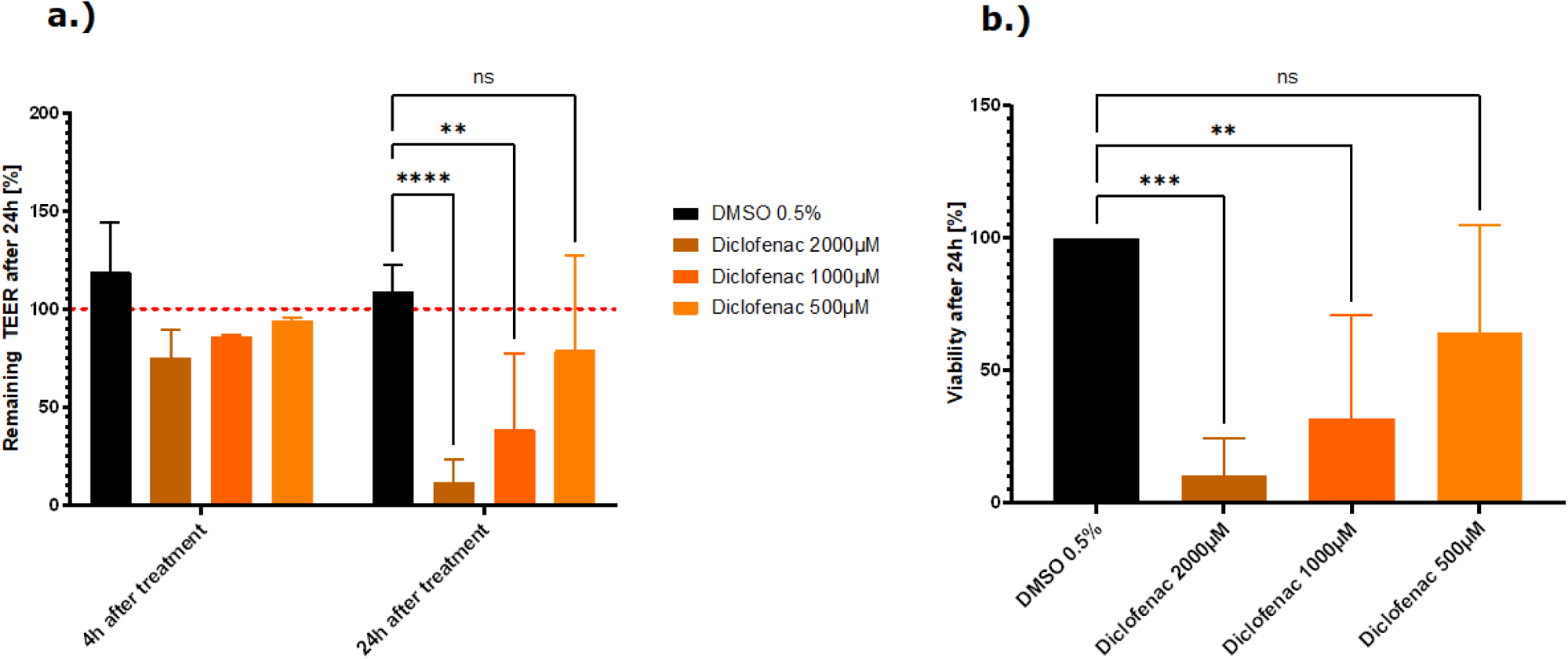
a.) Effect of the NSAID, diclofenac, on the barrier function represented as % TEER to the t0 of the treatment. Measured TEER values after treatment with the vehicle control DMSO and diclofenac. Shown are the mean remaining TEER values 4h and 24h after treatment. The red dotted line shows the normalized start TEER values of 100%. b.) Effect of the NSAID, diclofenac on the cell viability shown as % change to the DMSO control. Measured viability with the CellTiter Glo 3D kit, 24h after treatment with diclofenac The data are presented as means ± SD (n of controls=8, n of diclofenac= 3(2), statistical analysis of TEER values: **** p<0.0001, ** p=0.0078 and * p= 0.0241 by two-way ANOVA with Dunnett’s test for multiple comparisons to the control. For statistical analysis of Viability values: ****p < 0.0001 and ***p=0.0010 by one way ANOVA with Dunnett’s test for multiple comparisons to the control).

Treatment with phenylarsine oxide, patulin and homoharringtonine at the highest concentration, caused damage to the tight junctions resulting in a lower TEER value compared to the vehicle control, DMSO. Following the treatment with patulin and homoharringtonine, the TEER value decreased also at the lower concentrations tested (Figure 6a). From all the tight junction disruptors tested, patulin and phenylarsine oxide showed the strongest cytotoxicity with residual survival at approximately 0-20% at the highest concentration tested (100 and 5 µM, respectively) (Figure 6a). The TEER values decreased up to 80% after treatment of the cells with both compounds. After treatment the cells with the highest concentration of patulin and phenylarsine oxide the viability also decreased after 24h (Figure 6b). Sodium orthovanadate showed no toxic effect on the tight junctions or the vitality of the Caco-2 cells (Supplementary Figure 3)

**Figure 6:**
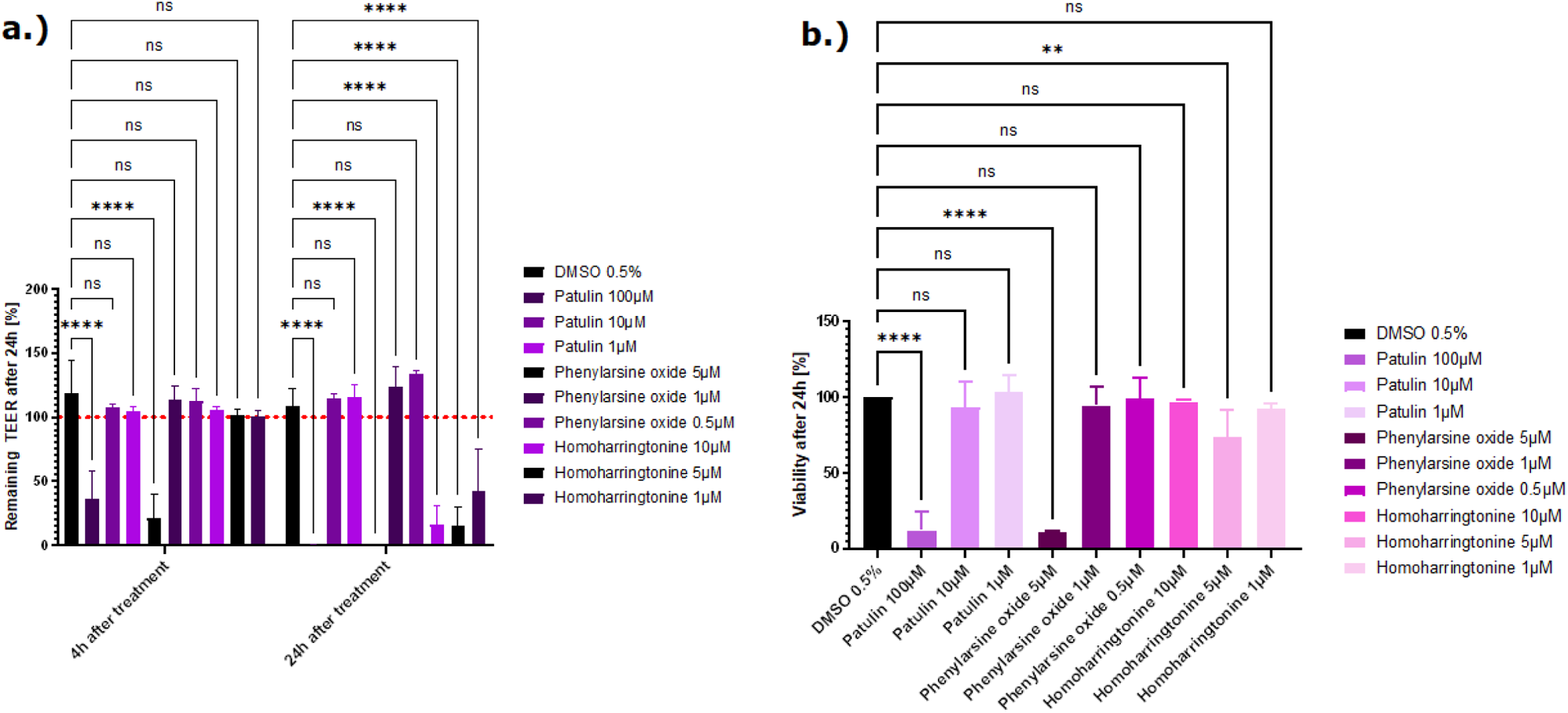
a.) Effect of three compounds from the tight junction damager group on the barrier function represented as % TEER to the t0 of the treatment. Measured TEER values after treatment with the vehicle control DMSO, patulin, phenylarsine oxide and homoharringtonine. Shown are the mean remaining TEER values 4h and 24h after treatment. The red dotted line shows the normalized start TEER values of 100%. b.) Effect of three compounds from the tight junction damager group on the cell viability shown as % change to the DMSO control. Measured viability with the CellTiter Glo 3D kit, 24h after treatment with patulin, phenylarsine oxide and homoharringtonine. The data are represented as means ± SD (n of controls = 8, n of patulin = 4, n of phenylarsine oxide = 2 and n of homoharringtonine = 3, statistical analysis of TEER values: **** p<0.0001 and ** p=0.0088 (phenylarsine oxide), 0.0070 (10µM homoharringtonine, 0.0074 (5µM homoharringtonine) by two-way ANOVA with Dunnett’s test for multiple comparisons to the control. For statistical analysis of Viability values: ****p < 0.0001 and **p=0.0072 by one way ANOVA with Dunnett’s test for multiple comparisons to the control).

Troglitazone and trovafloxacin showed no decrease of the TEER values and no decrease of viability. All Caco-2 tubes remained intact, and the cells were not affected by both liver toxic compounds (Figure 7).

**Figure 7:**
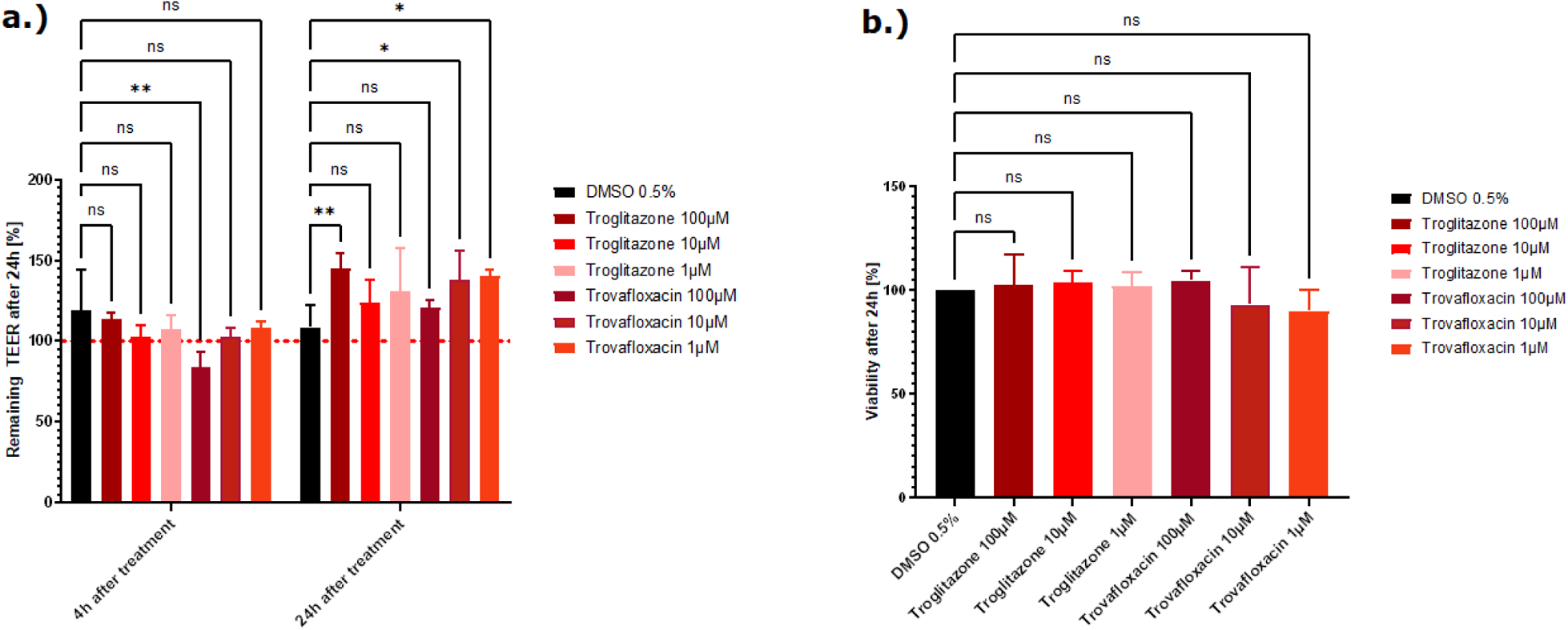
a.) Effect of the two liver toxins on the barrier function represented as % TEER to the t0 of the treatment. Measured TEER values after treatment with the vehicle control DMSO, troglitazone and trovafloxacin. Shown are the mean remaining TEER values 4h and 24h after treatment. The red dotted line shows the normalized start TEER values of 100%. b.) Effect of the two liver toxins on the cell viability shown as % change to the DMSO control. Measured viability with the CellTiter Glo 3D kit, 24h after treatment with troglitazone and trovafloxacin. The data are represented as means ± SD (n of controls = 8, n of troglitazone and trovafloxacin= 3, statistical analysis of TEER values: * p=0.0389 and ** p=0.0013 by two-way ANOVA with Dunnett’s test for multiple comparisons to the control. For statistical analysis of Viability values: one way ANOVA with Dunnett’s test for multiple comparisons to the control).

In this validation study most of the tested chemotherapeutic drugs did not show a relevant drop in the TEER value. Exposure to 5-FU, alosetron, irinotecan, sunitinib, sorafenib and carboplatin did not lead to a decrease of the TEER value nor to a decrease in viability (Supplementary Figure 5 and 6). Exposure to the highest concentration (300µM) of flavopiridol leads to a damage of the tight junctions with a subsequent decrease of the TEER value (approximately 60%) but the cells remained viable at this concentration (Supplementary Figure 4). Exposure to bortezomib leads to a decrease of the TEER value of a minimum of 70% at least in the highest concentration and bosutinib appeared to cause direct cytotoxicity and impacted the tight junctions, resulting in a simultaneous decrease in TEER and induction of cell death at 50 and 25µM (Figure 8a and 8b).

**Figure 8:**
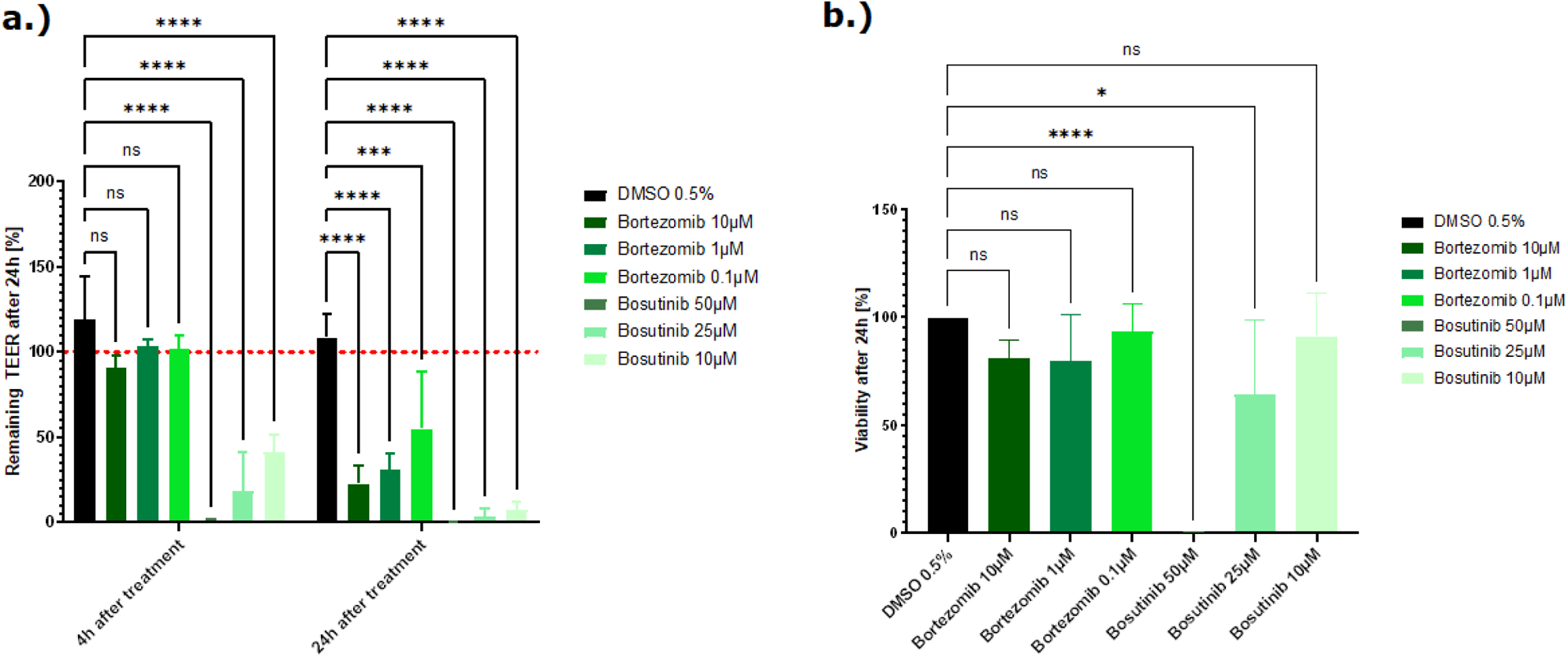
Effect of two chemotherauptic compounds on the barrier function represented as % TEER to the t0 of the treatment. Measured TEER values after treatment with the vehicle control DMSO, bortezomib and bosutinib. Shown are the mean remaining TEER values 4h and 24h after treatment. The red dotted line shows the normalized start TEER values of 100%. b.) Effect of two chemotherauptic compounds on the cell viability shown as % change to the DMSO control. Measured viability with the CellTiter Glo 3D kit, 24h after treatment with bortezomib and bosutinib. The data are represented as means ± SD (n of controls = 8, n of bortezomib and bosutinib= 3, statistical analysis of TEER values: **** p<0.0001, ***p =0.0005 (10µM bortezomib), *** p= 0.0003 (1µM bortezomib) and* p=0.0125 by two-way ANOVA with Dunnett’s test for multiple comparisons to the control. For statistical analysis of Viability values: **** p<0.0001 and * p= 0.0154 by one way ANOVA with Dunnett’s test for multiple comparisons to the control

The three compounds from the mixed group, loperamide, duloxetine and terfenadine, decreased at least at the highest concentration tested the TEER value and the viability decreased (Supplementary Figure 2). A summary of all results is shown in table 4.

**Table 4:**
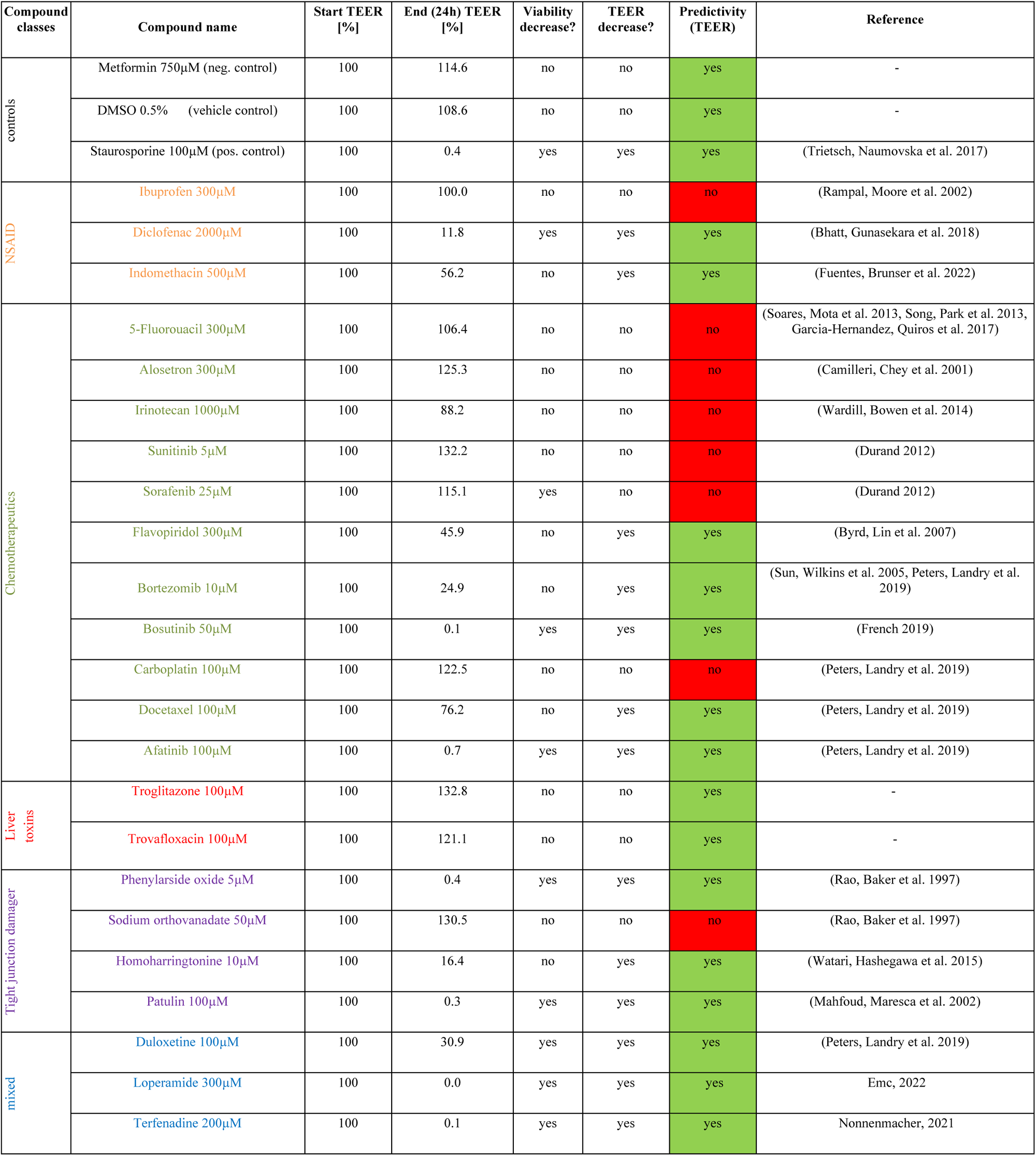
Overview of the TEER and viability results of the tested reference compounds (highest concentration tested). Shown are the % remaining start and end TEER values after treatment with the highest tested concentration of each compound.

In addition to these results, data from previous 2D Transwell studies were used to compare these with the results from this study to evaluate if the OrganoPlate® can help to better predict potential side effects of drug candidates on the intestinal barrier compared to the traditional Transwell models. For these studies Caco-2 cells were seeded onto semipermeable membranes of Transwell inserts which creates two compartments that mimic the intestinal barrier ((Awortwe, Fasinu et al. 2014), Figure 9). The cells were cultivated for 21days until they reached a confluent monolayer of >150 Ω*cm². The cells were then treated with test compounds and TEER values were determined after 24h. Overall, 9 test compounds (2 NSAID, 4 chemotherapeutics,1 liver toxin and 2 from a mixed group) were used in the Transwell experiments. Table 5 shows that 3 out of 9 compounds (loperamide, afatinib and terfenadine) lead to a decrease of the TEER values and the other 6 compounds (ibuprofen, indomethacin, 5-FU, alosetron, flavopiridol and troglitazone) did not influence the tight junctions and the TEER values remained stable.

**Figure 9:**
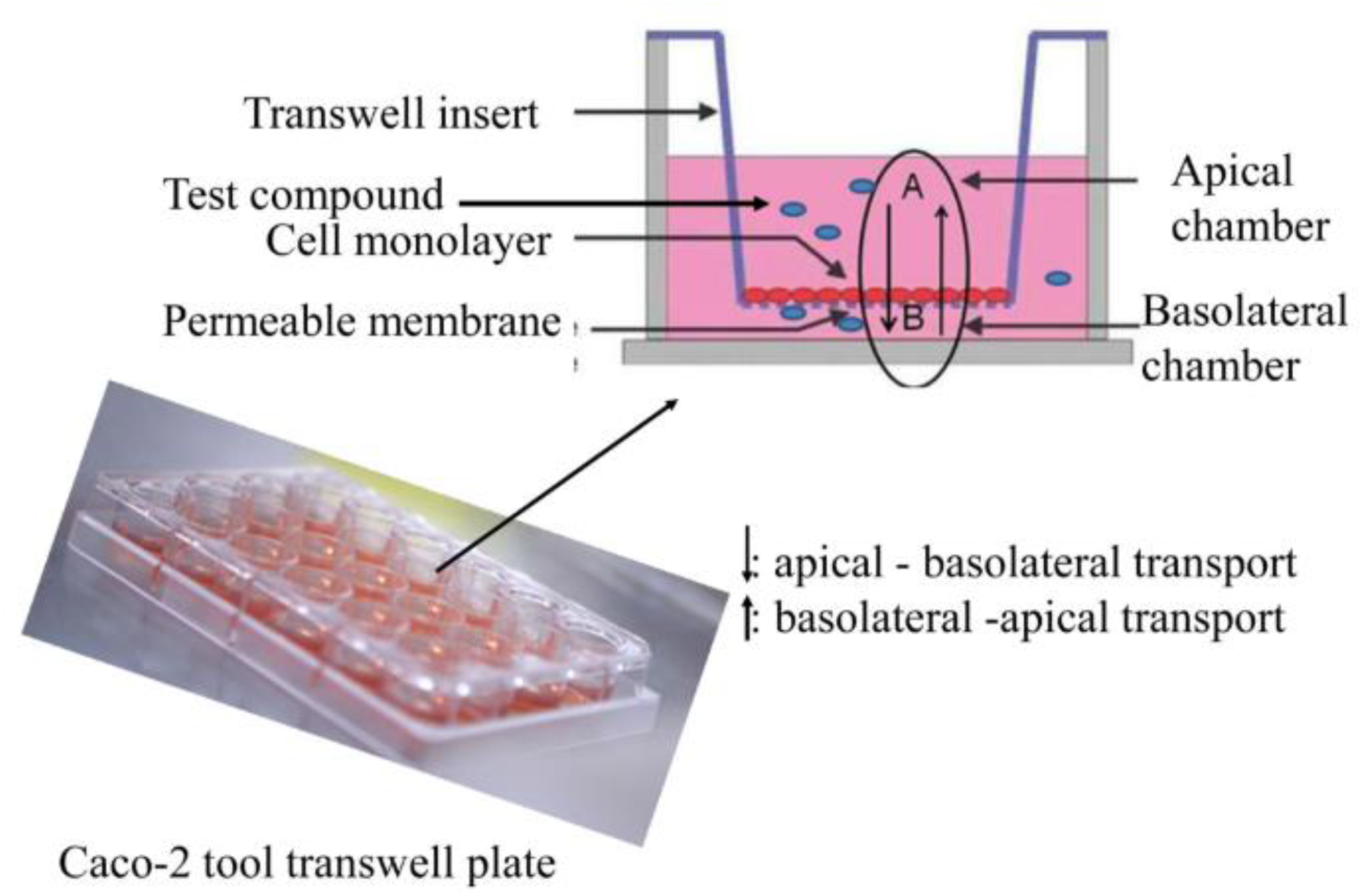
Cultivation of Caco-2 cells on permeable membranes of transwell inserts for transwell experiments ((Awortwe, Fasinu et al. 2014)

**Table 5:** Comparison of the TEER results from the OrganoPlate® and Transwell experiments and final evaluation of predictivity.

| Compound | OrganoPlate® (OP) |  |  | Transwell (TW) |  |  | OP vs. TW Predictivity |
| --- | --- | --- | --- | --- | --- | --- | --- |
|  | Start TEER [%] | End TEER [%] | TEER decrease? | Start TEER [%] | End TEER [%] | TEER decrease? |  |
| Metformin 750µM (neg. control) | 100 | **114.6** | no | 100 | **106.9** | no | OP = TW |
| DMSO 0.5% (vehicle control) | 100 | **108.6** | no | 100 | **103.1** | no | OP = TW |
| Staurosporine 10µM (pos. control TEER) | 100 | **1.7** | yes | 100 | **10.8** | yes | OP = TW |
| Ibuprofen 300µM | 100 | **100.0** | no | 100 | **108.5** | no | OP = TW |
| Diclofenac 2000µM | 100 | 11.8 | yes | NA | NA | NA | NA |
| Indomethacin 500µM | 100 | **56.2** | yes | 100 | **82.5** | no | OP > TW |
| 5-Fluorouracil 300µM | 100 | **106.4** | no | 100 | **90.1** | no | OP = TW |
| Alosetron 300µM | 100 | **125.3** | no | 100 | **131.5** | no | OP = TW |
| Irinotecan 1000µM | 100 | 88.2 | no | NA | NA | NA | NA |
| Sunitinib 5µM | 100 | 132.2 | no | NA | NA | NA | NA |
| Sorafenib 25µM | 100 | 115.1 | no | NA | NA | NA | NA |
| Flavopiridol 300µM | 100 | **45.9** | yes | 100 | **80.6** | no | OP > TW |
| Bortezomib 10µM | 100 | 24.9 | yes | NA | NA | NA | NA |
| Bosutinib 50µM | 100 | 0.1 | yes | NA | NA | NA | NA |
| Carboplatin 100µM | 100 | 122.5 | no | NA | NA | NA | NA |
| Docetaxel 100µM | 100 | 76.2 | yes | NA | NA | NA | NA |
| Afatinib 100µM | 100 | **0.7** | yes | 100 | **76.5** | yes | OP > TW |
| Troglitazone 100µM | 100 | **132.8** | no | 100 | **96.4** | no | OP = TW |
| Trovafloracin 100µM | 100 | 121.1 | no | NA | NA | NA | NA |
| Phenylarside oxide 5µM | 100 | 0.4 | yes | NA | NA | NA | NA |
| Sodium orthovanadate 50µM | 100 | 130.5 | no | NA | NA | NA | NA |
| Homoharringtonine 10µM | 100 | 16.4 | yes | NA | NA | NA | NA |
| Patulin 100µM | 100 | 0.3 | yes | NA | NA | NA | NA |
| Duloxetine 100µM | 100 | 30.9 | yes | NA | NA | NA | NA |
| Loperamide 300µM | 100 | **0.0** | yes | 100 | **5.9** | yes | OP = TW |
| Terfenadine 200µM | 100 | **0.1** | yes | 100 | **4.4** | yes | OP = TW |

## Discussion and conclusion

The OrganoPlate® 3-lane is a microfluidic, 3D cell culture platform which allows the differentiation and cultivation of Caco-2 cells in a tubular structure within 6 days. The medium flow, the cells being grown as 3D structures against a collagen layer and the fast differentiation of Caco-2 cells to small intestine-like enterocytes highlight the uniqueness of this model compared to commercially available 2D Transwell models. The OrganoTEER allows sensitive, real-time interrogation of compound effects on the intestinal barrier integrity in a throughput suitable for early preclinical safety assessment.

Overall, 23 reference compounds were used to evaluate the applicability of the OrganoPlate® (with Caco-2 cells) in combination with the OrganoTEER to determine toxic effects on the intestinal barrier. However, not all tested compounds showed damage to the tight junctions and thus a reduction of the TEER value.15 out 23 showed, at least the highest concentration tested, the expected outcome which means either a decrease in TEER value in parallel to a decreased viability after the treatment, a decrease in TEER value without a reduction of the viability, or a stable TEER value and no viability loss (Table 5).

The best predictivity was shown for the NSAIDs, the tight junction damager and the compounds which induce GI adverse effects in vivo (in each class 2 out of 3 compounds decreased the TEER value). For example, we were able to show that TEER values were reduced after treatment with the NSAID diclofenac, which indicates increased intestinal permeability. Consistent with a previous report, diclofenac can reduce TEER values, which indicates damaged tight junction connections and an increased intestinal permeability (Bhatt, Gunasekara et al. 2018)

Chemotherapeutic compounds are known to induce multiple adverse effects, including impacting the GI system, such as nausea, vomiting (Amjad, Chidharla et al. 2023) or diarrhea, pain and constipation (Lee, Ryan et al. 2014, McQuade, Stojanovska et al. 2016, Forsgård, Marrachelli et al. 2017). Diarrhea is often correlated with intestinal barrier damage and in 2019 Peters et al. published a method which uses a human GI microtissue and TEER for the investigation of the tightness of the intestinal barrier and the prediction of diarrhea. They showed that drugs with a very high probability of causing diarrhea in the clinic, like afatinib and idarubicin, with respectively 96% and 73% diarrhea incidence, correlate with a decrease in the TEER value and a disruption of the barrier (Peters, Landry et al. 2019). Our experiments showed the same results for afatinib, with TEER values decreased, indicating damage to the tight junctions, resulting in increased intestinal permeability, which is an indication of diarrhea. Nevertheless, the viability of the cells was also reduced after treatment with the two highest concentrations, suggesting that this effect in TEER maybe driven by direct cytotoxicity.

Our data show that the OrganoPlate® is only partially able to detect the damage to the intestinal barrier with chemotherapeutics. Only 5 of the 11 chemotherapeutics led to a reduction in TEER values. No effect on the intestinal barrier was observed with the other chemotherapeutics tested, in either the OrganoPlate or the Transwell systems. This can have different causes. For example, low predictivity in this context may be due to the treatment time. Chemotherapeutics have been developed to treat cancer cells, which usually have a very rapid division rate. In order to effectively treat this rapid proliferation, which can take several weeks to months, depending on the type and severity of the cancer, it is necessary to treat the cells several times. For example, colon cancer patients usually receive 4-6 months of chemotherapy (Sgouros, Aravantinos et al. 2015) which also increases the potential toxicity due to the long treatment with chemotherapeutics. Besides, some compounds only become toxic through metabolism in the liver and thus do not show direct toxicity in the intestinal cells. Irinotecan, for example, is metabolized to a toxic metabolite, SN-38, and the exposure to this active metabolite is correlated to diarrhea (Sun, Zhu et al. 2020). It is clear in the two GI in vitro models assessed here such metabolism is missing and therefore no direct toxicity is to be expected.

To demonstrate that the system does not produce false positives, specific compounds which induce liver injury, trovafloxacin and troglitazone, (Graham, Green et al. 2003) were also tested. Both compounds did not affect the tight junctions in the validation experiments nor decreased the viability of the Caco-2 cells (Figure 6 c & d), which supports the ability to predict correctly GI non-toxic compounds. No GI related toxicities have been reported for troglitazone (Klopotek, Hirche et al. 2006).

Phenylarsine oxide, patulin, sodium orthovanadate and homoharringtonine are all known to disrupt the tight junctions by either an inhibition of protein tyrosine phosphatase, a key regulator of intestinal epithelial barrier function (Rao, Baker et al. 1997, Mahfoud, Maresca et al. 2002) or by a downregulation of the claudine 3 and 4 expression. A disturbed localization of claudine 3 and 4 impairs the barrier formation (Watari, Hashegawa et al. 2015) and hence are essential for the function on the intestine. Clinically patulin is known to induce intestinal epithelial cell degeneration, ulceration, hemorrhages, and inflammation (Mahfoud, 2002) and leads to phosphorylation of claudin-4 and occludin which causes degradation of ZO-1 protein (Kawauchiya, Takumi et al. 2011) and can lead to a damage of tight junctions.

In our experiments phenylarsine, patulin and homoharringtonine decreased the TEER values in the OrganoPlate®, which indicated a damaged barrier. No data on these three substances could be collected for the 2D Transwell model to date.

The comparison of the results from the Transwell experiments and the OrganoPlate® experiments shows that the OrganoPlate® is more reliable in its ability to predict GI toxicity. The traditional Transwell was at least as predictive for intestinal barrier damage as the OrganoPlate® for 3 of the tested compounds. However, it is clear from our data that the Transwell model is not a suitable option for most of the other substances tested here, as it was unable to predict the toxic properties of the compounds on the intestinal barrier. In addition, Indomethacin, flavopiridol and afatinib led to a stronger decrease in the TEER value in the OrganoPlate® compared to the Transwell system. After using the Transwell system, these compounds would have been classified as non-toxic, whereas the OrganoPlate® was able to demonstrate clear damage to the intestinal barrier.

Overall, the TEER measurement is a reliable and widely accepted quantitative technique to identify the integrity of tight junctions of cell culture barriers (Srinivasan, Kolli et al. 2015). The present findings demonstrate that the measurement of TEER with the OrganoPlate®, can be used for the early safety prediction of drug-induced damage on the intestinal barrier. The OrganoPlate® includes several features, which are important for early pre-clinical drug screening (a) 3D tubular structure of cells which mimics the intestinal system itself, (b) sufficient throughput (40chips), (c) short pre-cultivation time compared to cell models on Transwell inserts (21days pre-cultivation) and (d) a functional endpoint which enable the distinguishment between drug-induced damage on the intestinal barrier (tight junctions) and cytotoxicity of the enterocytes. The data reported here helps the qualification of the OrganoPlate® as a routine test system for the early prediction of drug-induced GI toxicity. Further investigations are still required to improve the predictive outcome, for example by adding an additional cell type(s) to generate a co-culture platform (e.g., with HT29-MTX cells to simulate the mucus barrier) or by seeding iPSC derived gut organoids into the OrganoPlate® to mimic all the relevant cell types from the intestine. It is also advisable to test the metabolites of some of the compounds tested as the metabolism step in the liver can thus be indirectly considered.

## 1 Conflict of Interest

The authors declare that the research was conducted in the absence of any commercial or financial relationships that could be construed as a potential conflict of interest. KK, DK, WS are employees of Mimetas B.V., which markets OrganoPlate, OrganoTEER, and OrganoFlow, and holds the registered trademarks OrganoPlate, OrganoTEER, and OrganoFlow.

## 2 Author Contributions

The authors confirm contribution to the paper as follows: study conception and design: S.H. and P.H.; data collection: S. H. and I.K.; analysis and interpretation of results: S. H., P.H. and I.K.; draft manuscript preparation: S. H., P.H.; review of draft paper: P.H., W.S., D.K. and K.K.; approval of final version of the manuscript: S.H., P.H., I.K., W.S., D.K., K.K.

## 3 Funding

This work was supported by Mimetas.

## 4 Acknowledgments

The authors thank the following colleagues for their kind contribution to this work: A. Augustin and J. Langer.

## 6 Supplementary Material

Additional data evaluations are available in the supplementary material file.

## 7 Data Availability Statement

Datasets are available on request: The raw data supporting the conclusions of this article will be made available by the authors, without undue reservation.

