## Supplementary data for "Validation of an MPS based intestinal cell culture model for the evaluation of drug-induced toxicity"

### 1 Supplementary Data

#### 1.1 Supplementary Figures

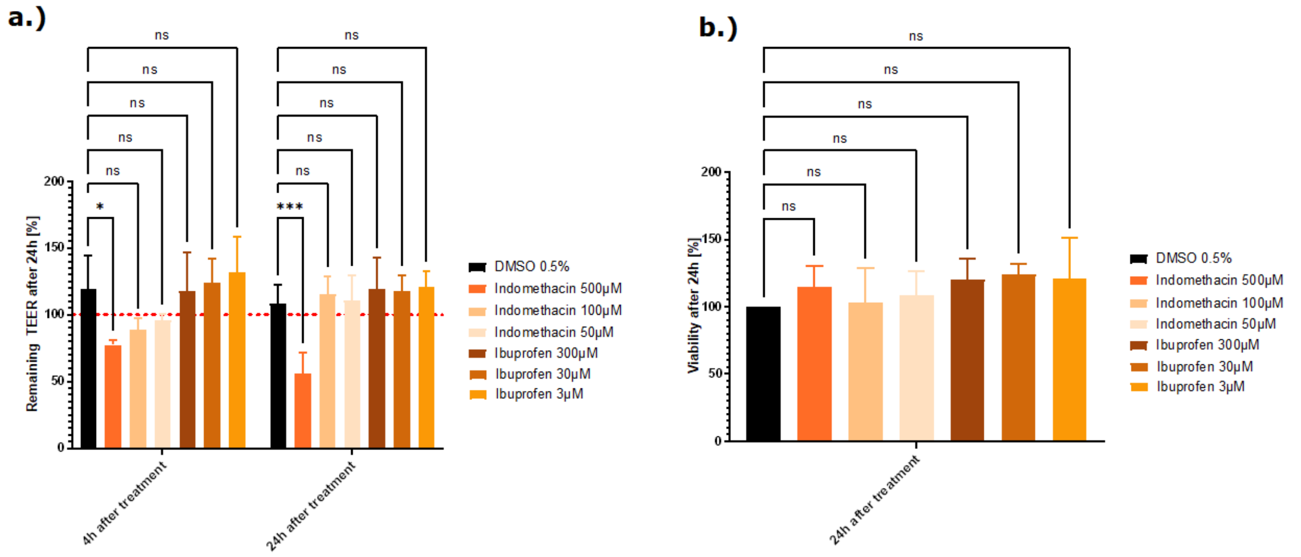


**Supplementary Figure 1: a.) Effect of two NSAID, indomethacin and ibuprofen, on the barrier function represented as % TEER to the t0 of the treatment. Measured TEER values after treatment with the vehicle control DMSO, the compounds indomethacin and ibuprofen. Shown are the mean remaining TEER values 4h and 24h after treatment. The red dotted line shows the normalized start TEER values of 100%. b.) Effect of two NSAID, indomethacin and ibuprofen on the cell viability shown as % change to the DMSO control. Measured viability with the CellTiter Glo 3D kit, 24h after treatment with Effect of the NSAID, diclofenac, on the barrier function represented as % TEER to the t0 of the treatment.indomethacin and ibuprofen = 3, statistical analysis of TEER values: * p=0.0102 (500µM indomethacin after 4h) and * p=0.0374 (500µM indomethacin after 24h) by two-way ANOVA with Dunnett's test for multiple comparisons to the control. For statistical analysis of Viability values: one way ANOVA with Dunnett's test for multiple comparisons to the control).**

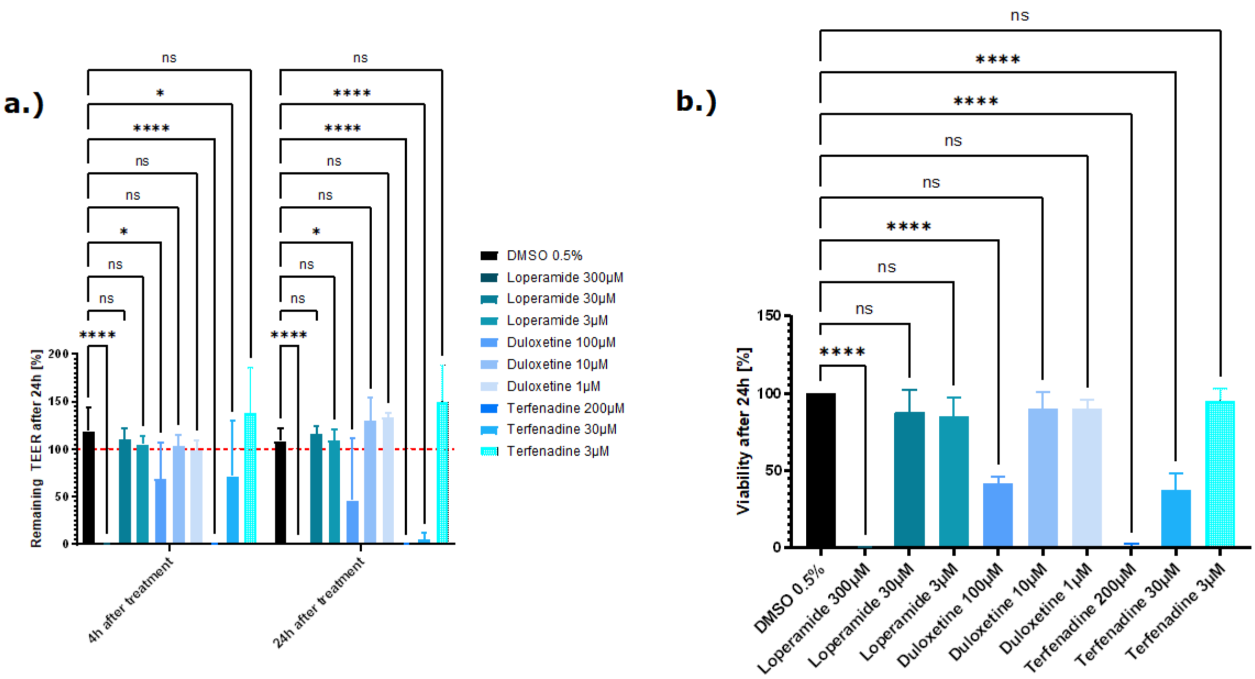


**Supplementary Figure 2: a.) Effect of the three compounds from the mixed group on the barrier function represented as % TEER to the t0 of the treatment. Measured TEER values after treatment with the vehicle control DMSO, the compounds loperamide, duloxetine and terfenadine. Shown are the mean remaining TEER values 4h and 24h after treatment. The red dotted line shows the normalized start TEER values of 100%. b.) Effect of the three compounds from the mixed group on the cell viability shown as % change to the DMSO control. Measured viability with the CellTiter Glo 3D kit, 24h after treatment with loperamide, duloxetine and terfenadine. The data are represented as means ± SD (n of controls = 8, n of loperamide = 4, n of duloxetine = 3 and n of terfenadine = 3, statistical analysis of TEER values: **** $p<0.0001$ and * $p=0.0160$ by mixed effects analysis with Dunnett's test for multiple comparisons to the control. For statistical analysis of Viability values: **** $p<0.0001$ by one way ANOVA with Dunnett's test for multiple comparisons to the control).**

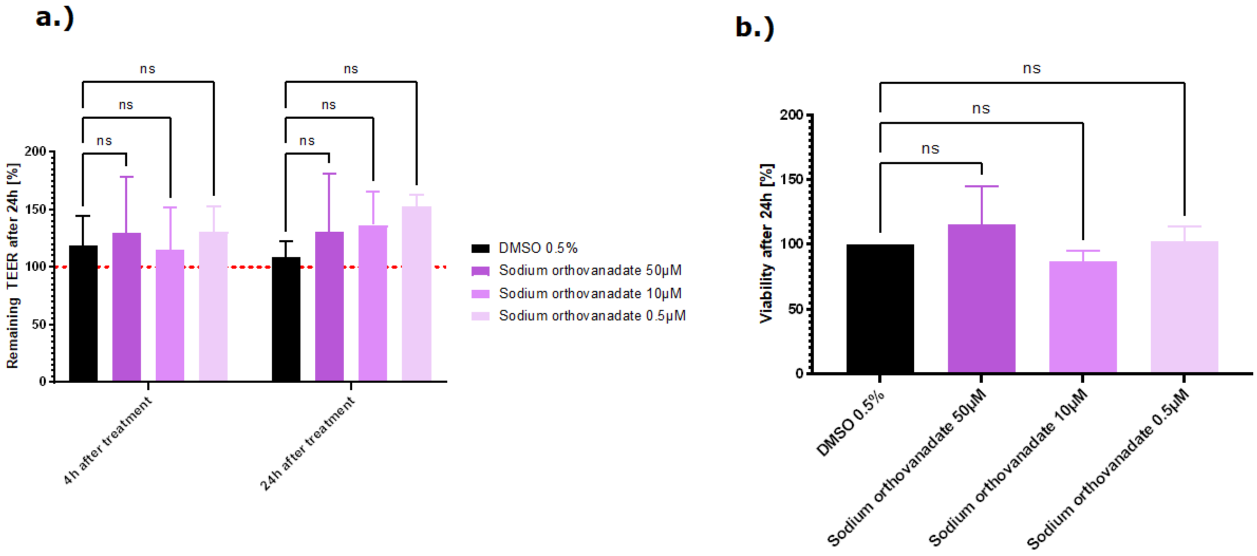


**Supplementary Figure 3: a.) Effect of sodium orthovanadate (tight junction damager) on the barrier function represented as % TEER to the t0 of the treatment. Measured TEER values after treatment with the vehicle control DMSO and the compound sodium orthovanadate. Shown are the mean remaining TEER values 4h and 24h after treatment. The red dotted line shows the normalized start TEER values of 100%. b.) Effect of sodium orthovanadate (tight junction damager) from the mixed group on the cell viability shown as % change to the DMSO control. Measured viability with the CellTiter Glo 3D kit, 24h after treatment with sodium orthovanadate. The data are represented as means ± SD (n of controls = 8, n of sodium orthovanadate= 3, statistical analysis of TEER values: ** p=0.0038 by two-way ANOVA with Dunnett's test for multiple comparisons to the control. For statistical analysis of Viability values: one way ANOVA with Dunnett's test for multiple comparisons to the control).**

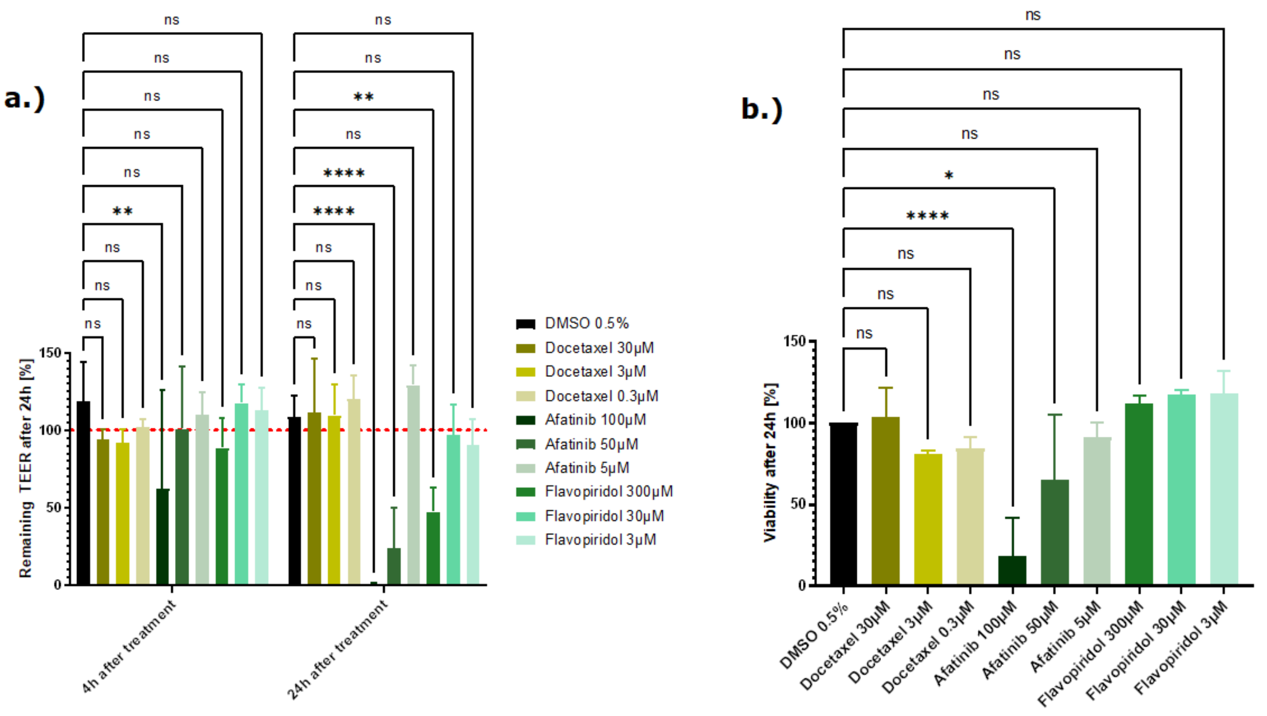


**Supplementary figure 4: a.) Effect of three chemotherapeutic compounds, docetaxel, afatinib and flavopiridol, on the barrier function represented as % TEER to the t0 of the treatment. Measured TEER values after treatment with the vehicle control DMSO, the compounds docetaxel, afatinib and flavopiridol. Shown are the mean remaining TEER values 4h and 24h after treatment. The red dotted line shows the normalized start TEER values of 100%. b.) Effect of three chemotherapeutic compounds, docetaxel, afatinib and flavopiridol, on the cell viability shown as % change to the DMSO control. Measured viability with the CellTiter Glo 3D kit, 24h after treatment with docetaxel, afatinib and flavopiridol. The data are represented as means ± SD (n of controls = 8, n of docetaxel, afatinib and flavopiridol = 3, statistical analysis of TEER values: **** p<0.0001 and * p=0.0253 by two-way ANOVA with Dunnett's test for multiple comparisons to the control. For statistical analysis of Viability values: **** p<0.0001 and *p=0.0339 by one way ANOVA with Dunnett's test for multiple comparisons to the control).**

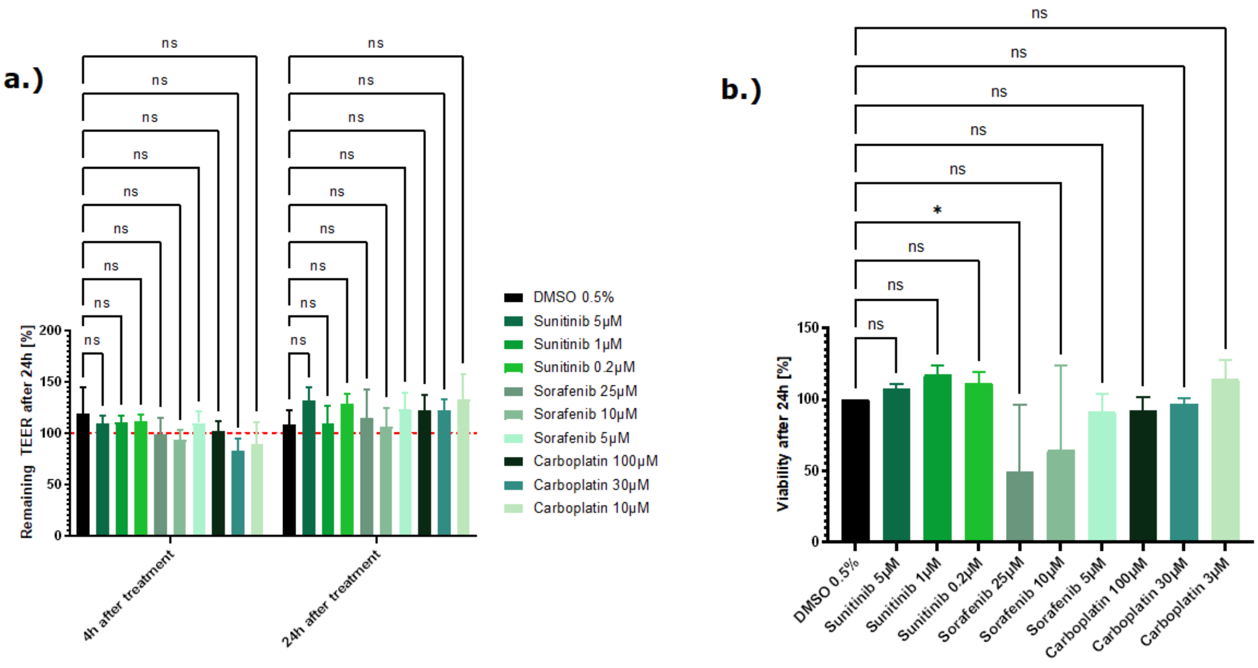


**Supplementary figure 5: a.) Effect of three chemotherapeutic compounds, sunitinib, sorafenib and carboplatin, on the barrier function represented as % TEER to the t0 of the treatment. Measured TEER values after treatment with the vehicle control DMSO, the compounds sunitinib, sorafenib and carboplatin. Shown are the mean remaining TEER values 4h and 24h after treatment. The red dotted line shows the normalized start TEER values of 100%. b.) Effect of three chemotherapeutic compounds, sunitinib, sorafenib and carboplatin, on the cell viability shown as % change to the DMSO control. Measured viability with the CellTiter Glo 3D kit, 24h after treatment with sunitinib, sorafenib and carboplatin. The data are represented as means ± SD (n of controls = 8, n of sunitinib and sorafenib = 3 and n of carboplatin = 2, statistical analysis of TEER values: two-way ANOVA with Dunnett's test for multiple comparisons to the control. For statistical analysis of Viability values: *p=0.0365 by one way ANOVA with Dunnett's test for multiple comparisons to the control).**

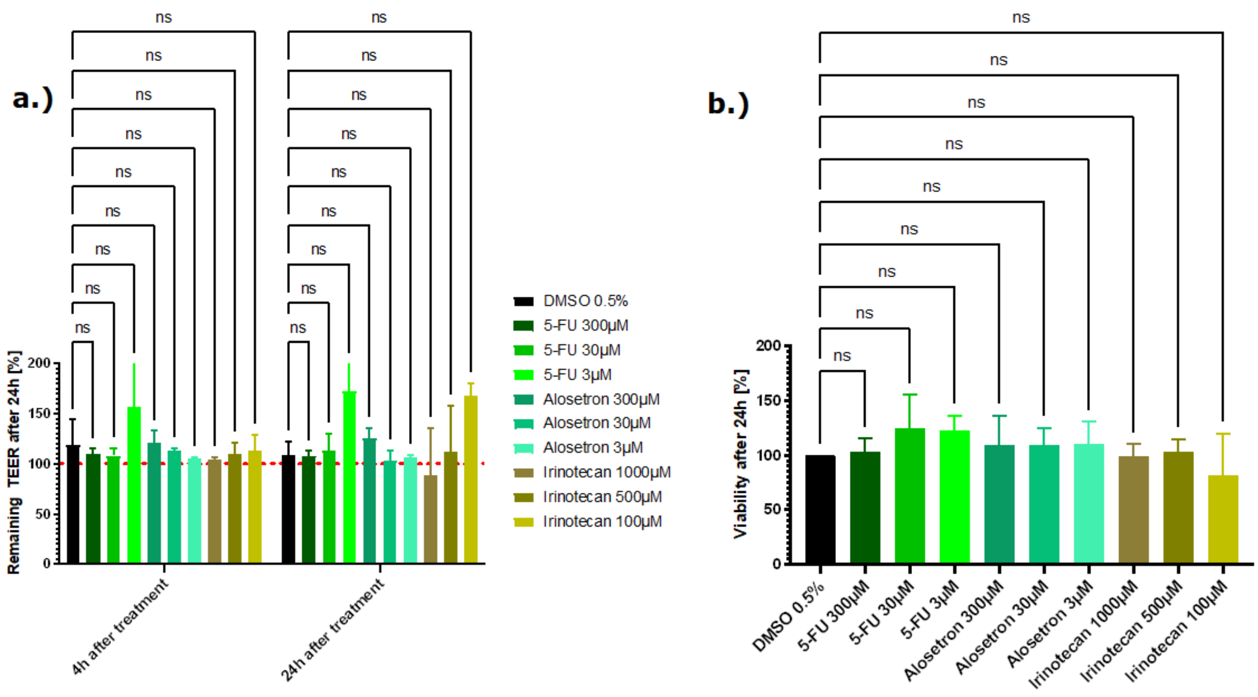


**Supplementary figure 6: a.) Effect of three chemotherapeutic compounds, 5-FU, alosetron and irinotecan, on the barrier function represented as % TEER to the t0 of the treatment. Measured TEER values after treatment with the vehicle control DMSO, the compounds 5-FU, alosetron and irinotecan. Shown are the mean remaining TEER values 4h and 24h after treatment. The red dotted line shows the normalized start TEER values of 100%. b.) Effect of three chemotherapeutic compounds, 5-FU, alosetron and irinotecan, on the cell viability shown as % change to the DMSO control. Measured viability with the CellTiter Glo 3D kit, 24h after treatment 5-FU, alosetron and irinotecan. The data are represented as means ± SD (n of controls = 8, n of 5-FU and irinotecan= 3 and n of alosetron = 2, statistical analysis of TEER values: * p= 0.0122 by two-way ANOVA with Dunnett's test for multiple comparisons to the control. For statistical analysis of Viability values: one way ANOVA with Dunnett's test for multiple comparisons to the control).**
